# Time-Resolved Cryo-EM Specimen Preparation with Single Millisecond Precision

**DOI:** 10.1101/2023.08.24.554704

**Authors:** Alejandra Montaño Romero, Calli Bonin, Edward C. Twomey

## Abstract

Molecular structures can be determined *in vitro* and *in situ* with cryo-electron microscopy (cryo-EM). Specimen preparation is a major obstacle in cryo-EM. Typical sample preparation is orders of magnitude slower than biological processes. Time-resolved cryo-EM (TR-cryo-EM) can capture short-lived states. Here, we present Cryo-EM Sample Preparation with light-Activated Molecules (C-SPAM), an open-source, photochemistry-coupled device for TR-cryo-EM with single millisecond resolution, tunable timescales, and broad biological applications.

---

Fundamental understanding of biological processes requires a description of their structural underpinnings. Through cryo-EM, three-dimensional (3D) biological structures *in vitro* and *in situ* can be reconstructed from molecular (∼10 Å) to atomic (∼1.2Å) resolution^1,2^. Samples are prepared for cryo-EM by mixing with a pipette, then depositing the mixture onto a cryo-EM grid. Excess sample or buffer is wicked away by filter paper, and the grid is plunge frozen into a cryogen. This procedure takes seconds to tens-of-seconds; in most cases, specimens are prepared on timescales that are orders of magnitude slower than the biological processes of interest; the overall molecular conformations that are fundamental to biology occur on the single millisecond (ms) to 1,000 ms timescale. Consequently, trying to correlate biological states observed from functional studies with conformational states reconstructed from cryo-EM is often challenging. Through TR-cryo-EM, non-equilibria states of molecules can be captured by rapidly preparing specimen across a temporal spectrum^3^. Generally, the approach of TR-cryo-EM is to activate the system at discreet time points prior to vitrification in the cryogen. Thus, with TR-cryo-EM, molecular conformations can be described as a function of time. The goal of the field has been to capture states on the milliseconds timescale to understand the short-lived states that are fundamental for the non-equilibria of biological processes^4^. Importantly, for drug development, TR-cryo-EM can be utilized with a background of small molecules to describe how they alter molecular conformational ensembles on a physiologically relevant timescale.

There are three principal approaches to TR-cryo-EM: i) on-grid mixing, ii) microfluidics, and iii) light-coupled (these techniques are reviewed in detail here^3^). We focused on light-coupled cryo-EM because of its cost effectiveness and adaptability. Light-coupled TR-cryo-EM can be applied to any system where photo-activatable ligands are available^5^ or to a system that is inherently photo-activatable (e.g., bacteriorhodopsin)^6^. Generally, this approach also only requires 2.5 to 3.0 μl of sample deposited on the cryo-EM grid, and thus offers an advantage to microfluidics where larger sample volumes are required to fill capillaries. In addition, the mixing approaches require arrays of different capillary chambers to facilitate different mixing times. Recently, light-coupled TR-cryo-EM devices have been made to analyze ligand-gated ion channels with high temporal resolution^7,8^. While these platforms are important developments, they have two drawbacks. First, the timescales are not readily adaptable, and second, single millisecond resolution, the theoretical limit of TR-cryo-EM^4,9^, was not achievable.

We report here the development of a new platform, Cryo-EM Sample Preparation with light-Activated Molecules (C-SPAM), which enables millisecond resolution light-coupled TR-cryo-EM and is adaptable to any timescale within single millisecond increments. Our idea was to construct a system where the light source and cryo-plunger are completely integrated together in a single embedded system. This allows us to control when exactly the light source could be coupled to the plunging of the sample. To achieve this, we initially built a light-emitting-diode (LED) into an already-existing open-access cryo-EM device^10,11^, but designed a new circuit board with an LED driver integrated into it (Extended Data Fig. 1). This controls each component of C-SPAM (Fig. 1a), all of which are either commercially available, have been previously designed and are open source, or are provided here and made open source (Extended Data Fig. 2). The main components include: i) a plunging solenoid, ii) a blotting solenoid, and iii) an LED (Fig. 1a). The plunging solenoid is connected to cryo-tweezers through a 3D-printed tweezer mount. The blotting solenoid sits on a 3D-printed support mount and is attached to a 3D-printed blotter mount that holds 25 mm filter paper. We constructed this initial version of C-SPAM with a 365 nm LED because the array of commercially available ligands that are photocleavable at 365 nm makes the device applicable to many biological questions (Table 1). This 365 nm LED can be exchanged for other commercially available light sources, making other systems such as optogenetics or thermal sensation available to TR-cryo-EM through C-SPAM (Table 1). The LED was constructed with a three-lens system, resulting in a 60 mm focal length, which defined where we placed the LED relative to the plunging path (Extended Data Fig. 3). This results in an irradiance of ∼1.3 W/cm^2^ at the cryo-grid, which is sufficient to uncage commercially available ligands at 365 nm^8^. While the heat generated from light activation has been a concern, higher powered LEDs than here generate minimal heating and do not affect freezing^7^.

**Fig. 1.**
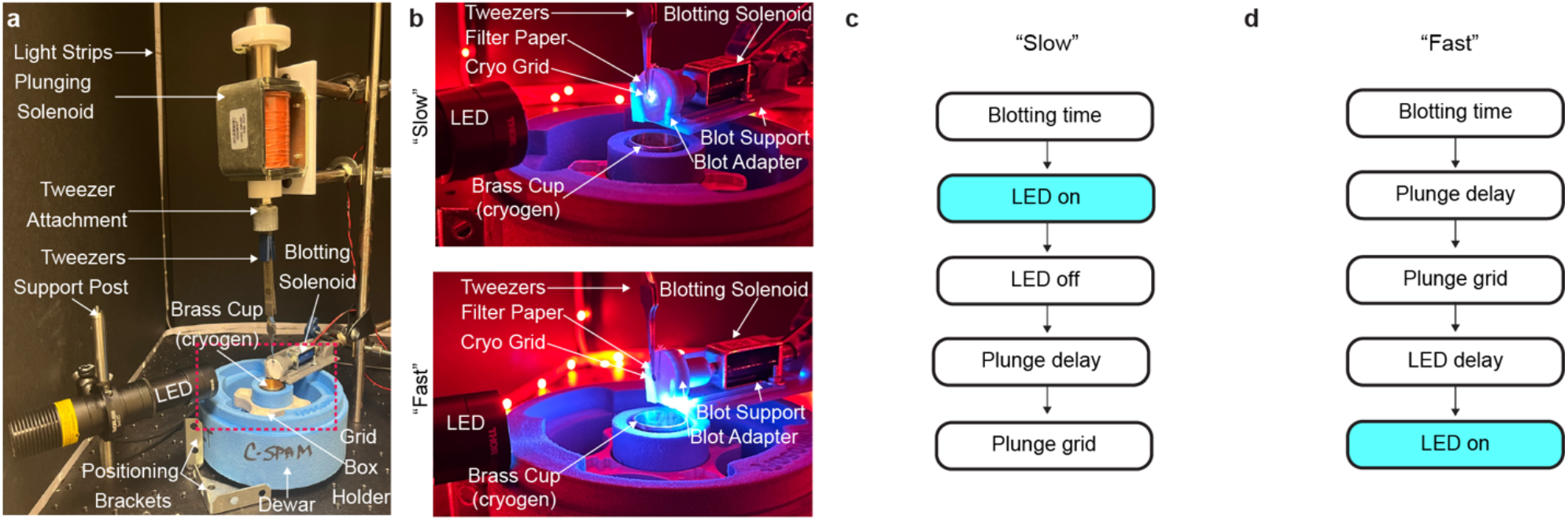
C-SPAM build and operation. **a**, Overall construction of C-SPAM. All components necessary for operation are labeled. **b**, Two operations are available with C-SPAM, “Slow” and “Fast”. The “Slow” operation has the LED at the top position, illuminating the cryo grid at its starting position. The “Fast” operation has the LED at the bottom position, illuminating the cryo grid directly above the brass ethane cup. **c-d**, Order of functions for “Slow” and “Fast” operations.

**Table 1.**
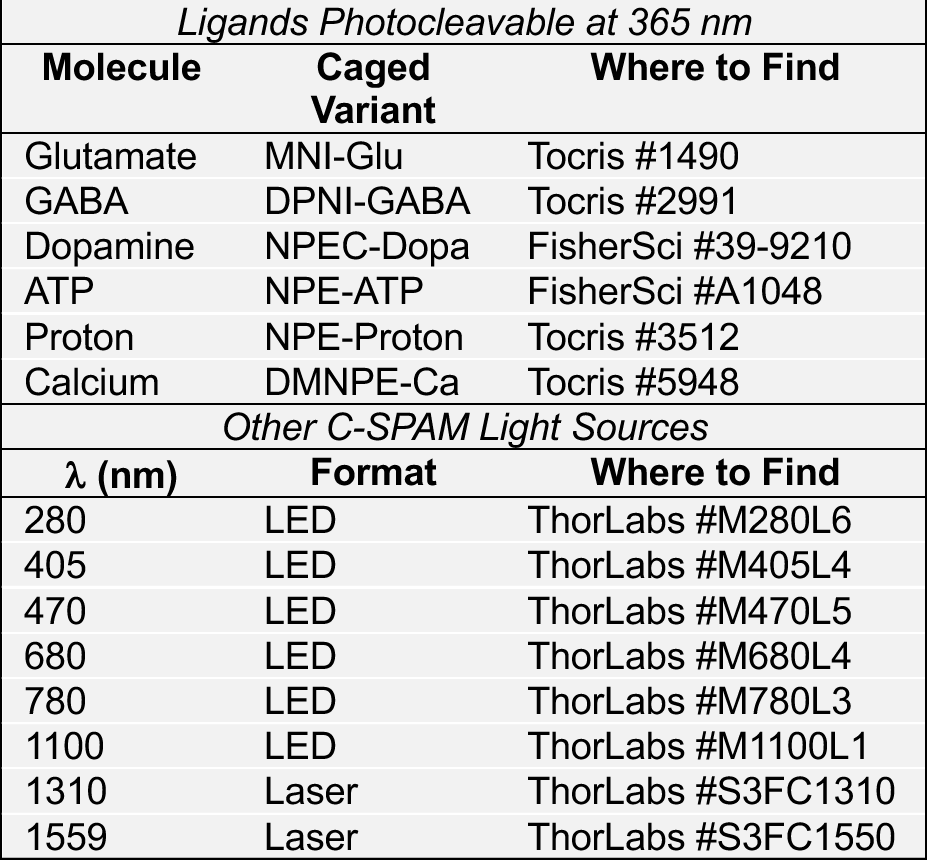
Uses and Adaptations of C-SPAM.

There are two positions that the LED can occupy depending on the desired temporal resolution, i) “Slow” and ii) “Fast” (Fig. 1b). The construction of the device allows for easy and convenient movement between these two positions. In the “Slow” position, the LED irradiates the sample for a defined period prior to plunging (Fig. 1c). The LED is positioned to where it is aligned on the cryo grid as it rests on the tweezers before plunging. This configuration enables TR-cryo-EM for time periods of 100 ms or slower. In the “Fast” position, the LED irradiates the sample during plunging before it enters the cryogen (Fig. 1d). Here, the LED is placed flush against the dewar, which results in the area just above the brass ethane cup being illuminated. In this position, a temporal resolution of 1-15 ms is achievable.

To reach rapid and adaptable temporal resolutions with C-SPAM, we used a standardization technique that we encoded into the software. For precise standardization, we used a slow-motion camera with a 960-fps setting (∼1.04 ms/frame). The recording started from the movement of the plunging solenoid, through LED powering, and ended once the grid was plunged into the ethane cup, representing vitrification (Fig. 2a). By accounting for the number of frames between each point of the process, we determined the time in milliseconds for the grid to descend from its starting point to the liquid ethane. We then defined an LED power delay that is defined as the amount of time (ΔT_1_) the LED waits to turn on once the plunging solenoid is activated (Fig. 2b). The powering of the LED activates the system, and the time between this and vitrification in the cryogen (ΔT_2_) is the temporal resolution (Fig. 2b). This total value of time is ΔT_1_ + ΔT_2_. Users of C-SPAM can input which temporal resolution their sample need to be prepared at, which is defined as LED delay = (ΔT_1_ + ΔT_2_) – resolution input (Fig. 2c). Altogether, this allows for tunable temporal resolution with single-millisecond precision.

**Fig. 2.**
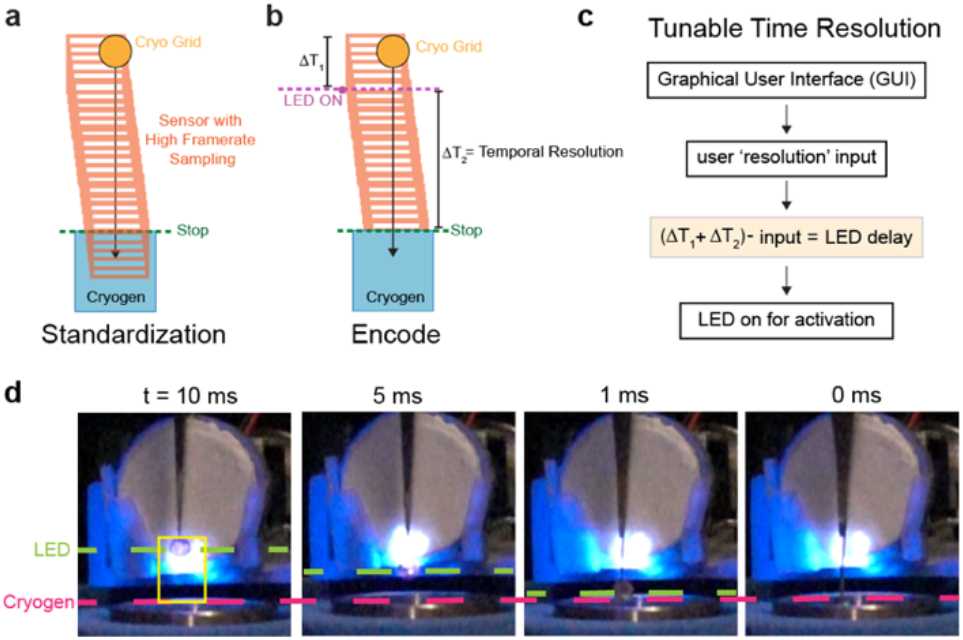
Temporal resolution standardization. **a**, Standardization of C-SPAM uses a high framerate sampling technique, considering the amount of time the cryo grid takes to descend into cryogen from its starting position. **b**, Encoding of C-SPAM time resolution defines descending time as (ΔT_1_ + ΔT_2_), where ΔT_1_ is time from start to LED on and ΔT_2_ is time from start of reaction to vitrification, representing temporal resolution. **c**, For tunability of time resolution, an LED delay is encoded in the software that is dependent on both the user resolution input and (ΔT_1_ + ΔT_2_). LED delay is defined as (ΔT_1_ + ΔT_2_) – input. **d**, Images of 10, 5, 1, and 0 ms temporal resolutions of C-SPAM.

To prepare samples with single-millisecond precision using C-SPAM, the user uses the graphical user interface (GUI) that allows for tunable times for blotting, resolution, and intensity of the LED (Extended Data Fig. 3). Once these parameters are determined for a specific biological system, the user begins by adding sample directly to a grid. As the program starts, the blotting solenoid moves forward to blot the grid, removing excess sample. Once finished, the blotting solenoid retracts and engages the plunging solenoid to begin moving the grid towards the cryogen. The LED turns on at a specified point determined by the selected time resolution, activating the reaction (Fig. 2b). For example, we show C-SPAM being utilized for TR-cryo-EM at 10 ms, 5 ms, and 1 ms (Fig. 2c, Extended Data Fig. 4). At each of these time points, through our standardization and encoding of the embedded system, the LED turns on at exactly the user-defined time resolution prior to freezing. If the user is using the “Slow” parameters, the process changes by irradiating the grid for the desired time following blotting (Fig. 1c). To verify that the system can prepare sample without irradiation damage, we collected single particle cryo-EM data on and generated a 3D reconstruction of apoferritin with C-SPAM (Extended Data Fig. 5).

C-SPAM enables investigation of conformational dynamics at physiological time scales with cryo-EM and is adaptable to many areas of biology. In addition, C-SPAM aims to address overarching barriers in TR-cryo-EM such as the requirements of financial capital, engineering expertise, and high sample volume. All components of this device are available and open source to spur community driven development of TR-cryo-EM.

## Methods

### C-SPAM Instrument

#### C-SPAM Build

The instrument was built using 3D prints outlined in the Rubenstein device^10,11^, along with new incorporations and designs as specified. All part names and their availability used in C-SPAM, circuit board schematics, and code can be found on https://github.com/twomeylab/c-spam.

The C-SPAM device is built on a breadboard (ThorLabs MB2424) that holds the plunging and blotting solenoids, along with the LED construction. The plunging solenoid (Summit Electronics HD8286-R-F – 24VDC) is supported by a 3D printed holder^11^ and contains a 3D printed piece containing magnets (6×3MM, purchased from Amazon, secured with epoxy) that sits on the solenoid head^11^. The 3D printed holder is mounted on a linear motion shaft as described^11^. This is essential for keeping the solenoid in place while de-energized. The plunging solenoid piston contains a 3D printed spacer^11^ followed by a 3D printed tweezer attachment designed in our lab (Extended Data Fig. 2). The tweezer attachment is compatible with Nanosoft’s cryo-tweezers (Nanosoft 17021002). Furthermore, the base of the blotting solenoid (Digikey 1528-1551-ND) is supported by a 3D printed support and is connected to a 3D printed blot adapter, both designed our lab (Extended Data Fig. 2). The LED (ThorLabs M365L3) is attached to an optical post (ThorLabs TR6) via a lens tube coupler (ThorLabs SM1TC) and has a three-lens design attached. The optical post is attached to a second optical post through a swivel clamp (ThorLabs SWC/M), which allows the height of the LED to be easily adjusted.

There are three coupled lens tubes. The first lens (ThorLabs ACL25416U-A) is held at the front of the SML10 lens tube, followed by the second lens (Thor Labs LB1596-A) held at the end of the SML03 lens tube, and finishing with the third lens (Thor Labs AC254-050-A-ML) held at the end of the SM1V05 lens tube (Extended Data Fig. 3). The LED design results in a 60 mm focal length. To measure irradiance, we used a USB Power Meter Thermal Sensor 0.19 -20 μm 10 W Max (ThorLabs PM16-425). All irradiance measurements were taken by placing the power meter 60 mm from the LED build and shining the LED at 100% via the C-SPAM GUI. All readings are using a 3 mm diameter and a flat top beam profile.

The C-SPAM build uses Nanosoft’s vitrification dewar (Nanosoft 21021005), along with its compatible brass ethane cup (Nanosoft 21031002) and grid box holder (Nanosoft 21021002) and is held in place by positioning brackets. All hardware components described are connected to the circuit board (Extended Data Fig. 1). The circuit board is integrated directly onto the Raspberry Pi 3 Model B+. The full C-SPAM device is powered by a 24V power supply.

#### C-SPAM Software

All scripts described are written in Python 3. There are three main scripts that makeup the C-SPAM software: i) CSPAMgui.py, ii) CSPAMpinlist.py, and iii) CSPAMfunctions.py. The Graphical User Interface (GUI) that is used is encoded by the CSPAMgui.py script. The GUI contains four parameters that can be user-defined: i) blotting time (ms), ii) plunge delay (ms), iii) resolution (ms) and iv) LED intensity (%). In addition, the GUI contains three buttons: i) Ready, ii) Start, and iii) Abort (Extended Data Fig. 3). The initial GUI was adapted from the Rubinstein device^11^. The Ready button will bring the blotting solenoid forward. The Start button will begin the full automated process described. The Abort button will stop all processes. The GUI also contains a checkbox of ‘Do Not Plunge’ to impede the plunging solenoid from being energized. Once the process is started by the user, the hardware components are called by the CSPAMpinlist.py script and the CSPAMfunctions.py module is imported. The automated process then begins and utilizes an if-else statement to differentiate between the two possible time scales. If the user-defined resolution input is ≤ 15 ms, the “Fast” process is started and the applyandplungeFAST function (imported from the CSPAMfunctions.py module) is called. If the resolution is ≥ 50 ms, the “Slow” process is started and the applyandplungeSLOW function is called.

### Standardization of Time Resolution

To standardize the time resolution achievable by C-SPAM, we used a high frame-rate sampling technique with a slow-motion camera. With C-SPAM set up in the “Fast” position, we took 960-fps (∼1.04 ms/frame) videos with a Sony RX0 II 1” (1.0-type) sensor ultra-compact camera. Videos were analyzed using Adobe Premiere Pro. We accounted for the number of frames it took for the grid to reach vitrification from its original position at the start of plunging. This frame number represents the total time the plunging to vitrification process takes, represented by (ΔT_1_ + ΔT_2_) (Fig. 2). We used this value to determine an LED delay, which is defined as the amount of time the LED waits to turn on once the plunging solenoid is activated. We encoded this value as part of the leddelay argument in the scripts. To have tunable time resolution with single-millisecond precision, we made a final LED Delay function that is dependent on both the frame value observed and the user-defined resolution input. The final LED Delay is defined as (ΔT_1_ + ΔT_2_) – resolution input.

### Operation

C-SPAM has two main operations, “Fast” and “Slow”. Both operations begin by the user bringing up the GUI and filling out the parameters. The user then adds 3 μl of sample to the cryo-grid. With the press of the Ready button, the blotting solenoid comes forward to blot excess sample off the grid. If using the “Fast” operation, upon pressing Start, blotting time will begin. Once finished, the blotting solenoid will retract, initiating the plunge delay time. This plunge delay time is essential to prevent crashing of the tweezers and blotting solenoid. Once the delay is finished, this activates the plunging solenoid to descend towards the cryogen.

With the plunging solenoid activated, this calls the LED delay previously described and will turn on the LED at the desired time resolution prior to vitrification. If using the “Slow” operation, the LED will turn on once the blotting has finished. The LED will remain on for the defined resolution time and the plunging solenoid will become activated once the LED turns off. The user can either replace or rotate the 25 mm Whatman paper between each grid.

### Cryo-EM

All data was collected on a 200 kV ThermoFisher Glacios equipped with a Falcon 4i camera using ThermoFisher EPU software. For the apoferritin collection, 3 μl 4 mg/ml mouse apoferritin was plunge frozen with C-SPAM in “Slow” mode after 4 s blotting with 25 mm Whatman #1 filter paper. Micrographs were collected at 1.17 Å/pixel with a 40 e^-^ /Å^2^ dose. All data were imported into Cryosparc with 40 frames and no upsampling. 154 micrographs were used for processing, and blob picking was used to generate templates for picking. Pickled particles were extracted with a 250 pixel box size. A subset of particles was used in ab initio reconstruction to generate an initial reference of apoferritin, which was refined to 3.55 Å with a group of 43,620 particles and octahedral symmetry applied. This reconstruction was used as a 3D reference in heterogeneous refinement. 8,945 particles from the best class were reconstructed to 3.26 Å with octahedral symmetry.

## Data Availability

All 3D printables, circuit designs, code, and scripts designed as part of this study are available at https://github.com/twomeylab/c-spam. All other data and meta data from this study are available from the corresponding author upon request.

## Acknowledgements

E.C.T. is supported by the Searle Scholars Program (Kinship Foundation, # 22098168) and the Diana Helis Henry Medical Research Foundation (#142548). Cryo-EM data was collected at the Beckman Center for Cryo-EM at Johns Hopkins. We thank Ye Ma and Taekjip Ha (Johns Hopkins) for assistance with the LED lens design, the Instrument Development Group (Johns Hopkins) for assistance and assembly of the circuit design, and James Berger (Johns Hopkins) and Duncan Sousa (Johns Hopkins) for helpful discussions. The apoferritin sample was a gift from the Berger lab at Johns Hopkins.

## Author Contributions

A.M.R. and E.C.T conceived the project, designed the project, and prepared the manuscript with assistance from all authors. A.M.R. and C.B. constructed the device, software, and collected all data. A.M.R. and C.B. prepared the cryo-EM samples and conducted the cryo-EM experiments. E.C.T. supervised all aspects of the study.

## Competing Interests

Johns Hopkins University has filed a provisional patent based on this work. The authors declare no competing interests.

**Extended Data Fig. 1.**
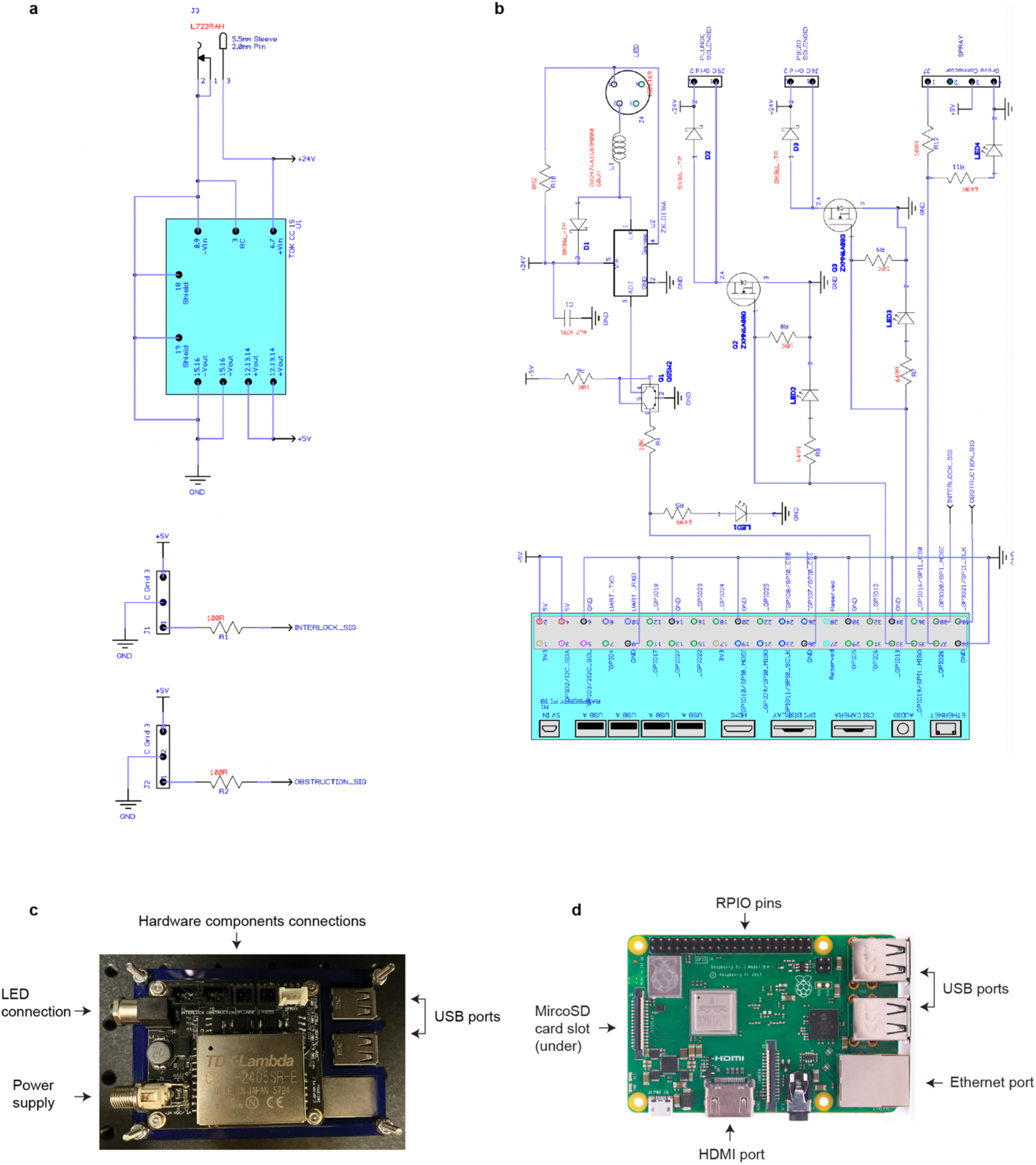
Circuit schematic and electrical components of C-SPAM. **a-b**, Circuit board schematic for incorporation onto Raspberry Pi Model 3 B+. **c**, Fully assembled circuit and Raspberry Pi Model 3 B+. Connections for LED, hardware components, USB ports, and power supply are shown. **d**, Overview of Raspberry Pi Model 3 B+.

**Extended Data Fig. 2.**
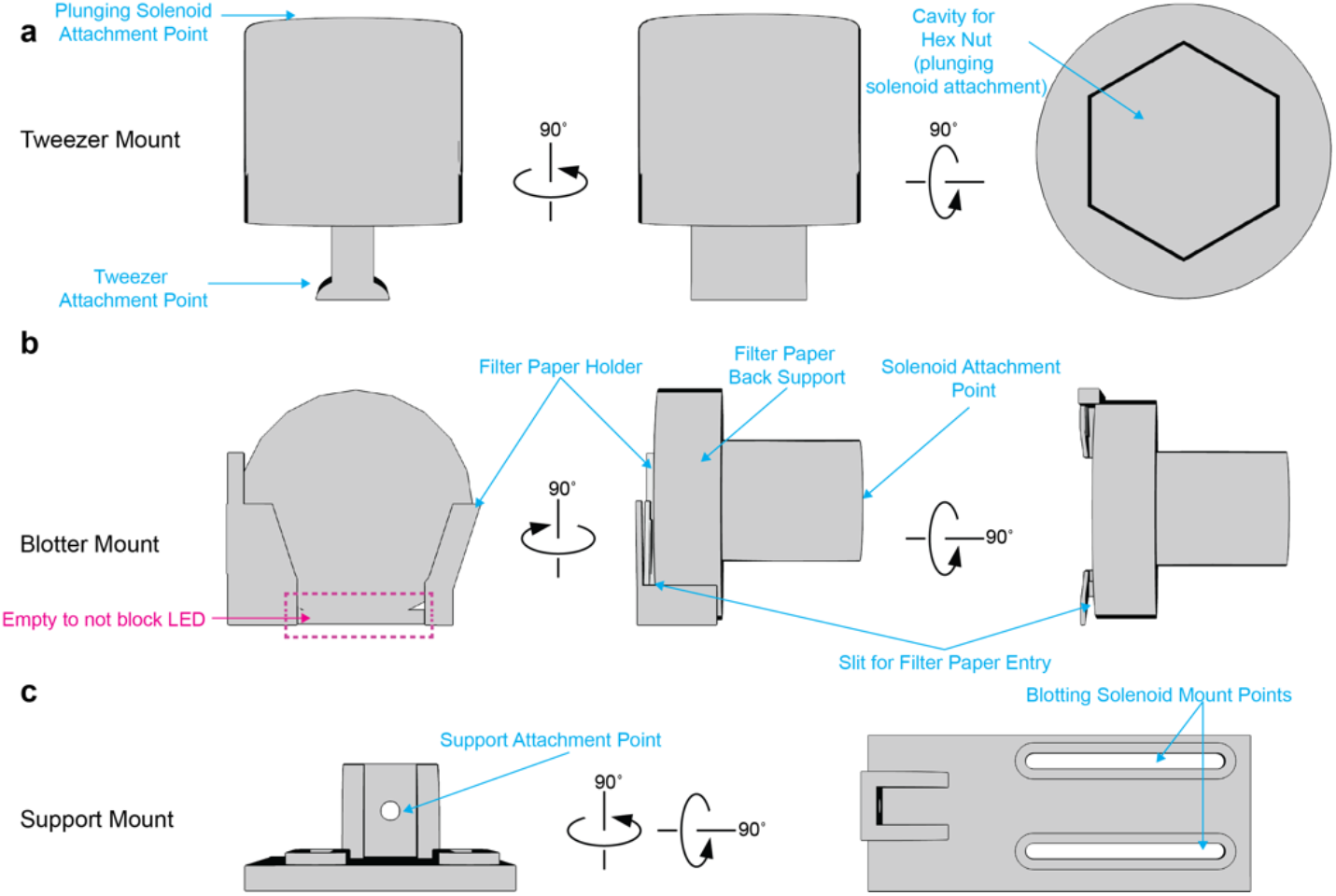
Custom 3D print designs for C-SPAM. **a**, Schematic of tweezer mount, showing all important attachment points. **b**, Schematic of blotter mount, showing placement points for 25mm filter paper and solenoid attachment. **c**, Schematic of support mount for the blotting solenoid.

**Extended Data Fig. 3.**
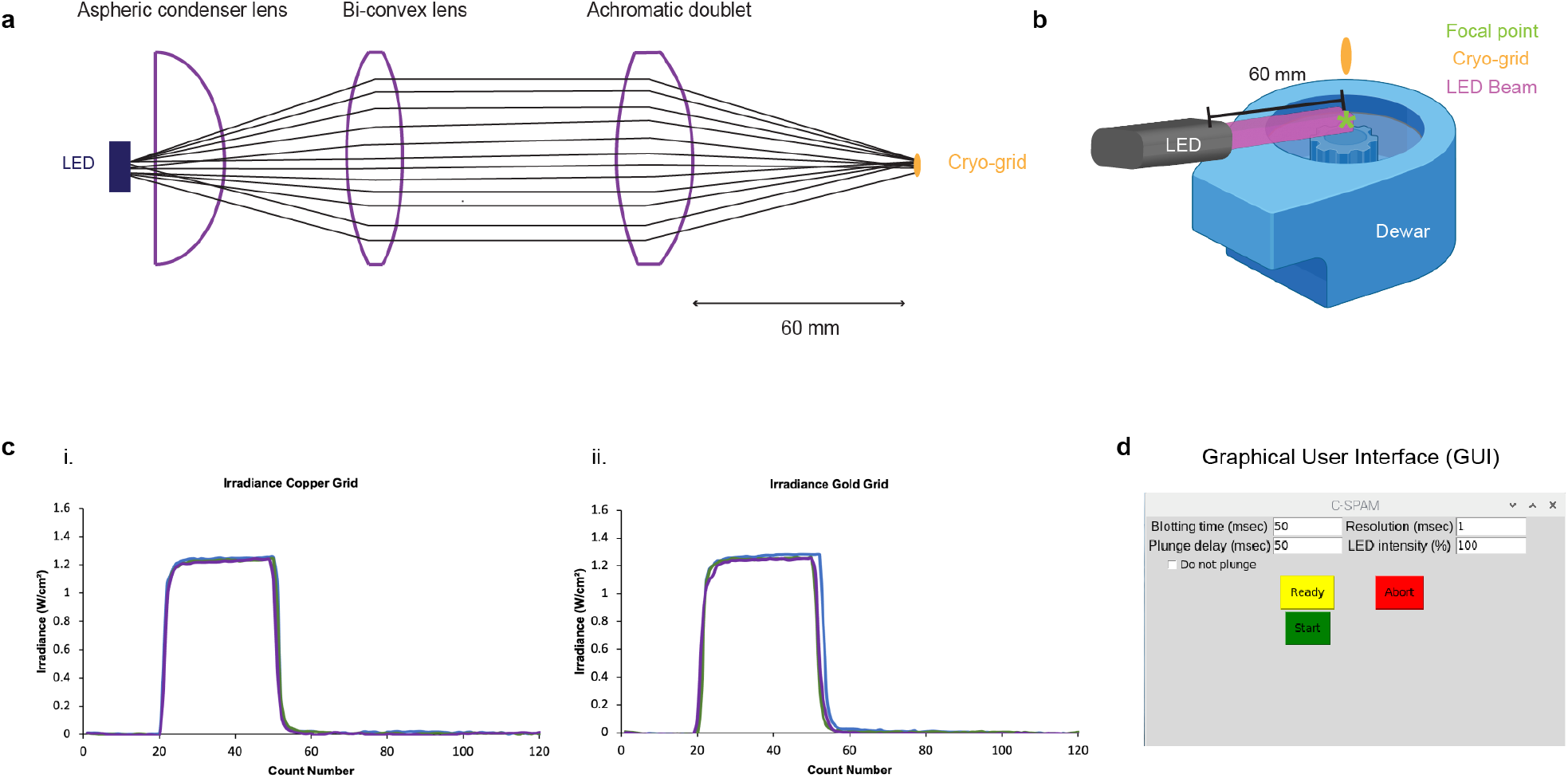
LED lens design, irradiance, and GUI. **a**, Diagram of LED three-lens design, depicting order and function of each lens. **b**, Overview of relationship between LED focal point and cryo grid. **c**, (i) Irradiance (W/cm^2^) of LED at 100% with copper grid present shows a maximum of ∼1.3 W/cm^2^, (ii) Irradiance (W/cm^2^) of LED at 100% with gold grid present shows same maximum. Each were run in triplicate. **d**, C-SPAM Graphical User Interface (GUI) showing all available tunable parameters.

**Extended Data Fig. 4.**
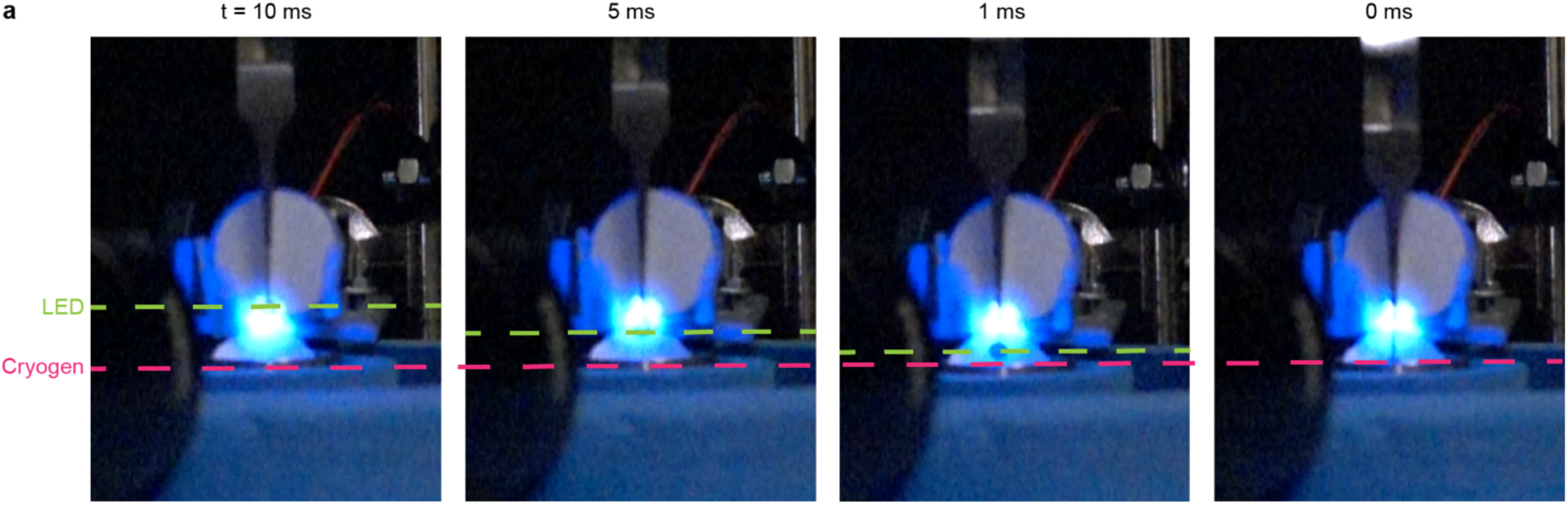
Temporal resolution frames. **a**, Frames representing three different temporal resolutions are shown: 10, 5, and 1 ms. 0 ms frame shows represents vitrification. Green dashed line shows activation point and magenta dashed line shows vitrification point. Whatman filter paper was placed in brass ethane cup to show full LED beam diameter.

**Extended Data Fig. 5.**
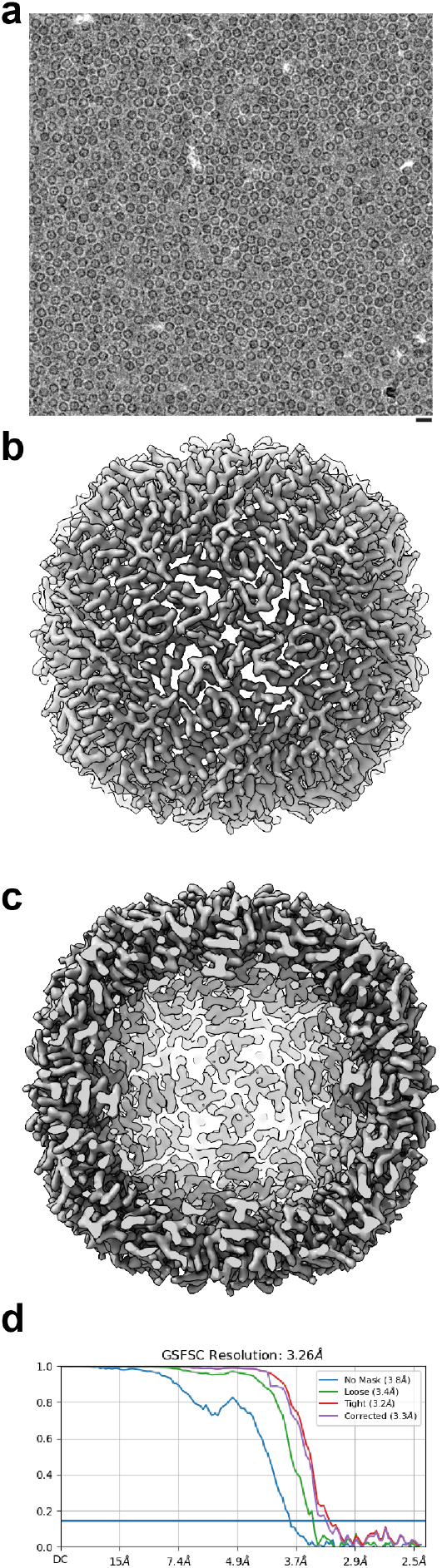
Apoferritin reconstruction using C-SPAM. **a**, Representative apoferritin micrograph. 20 nm scale bar. **b-d**, Reconstruction of apoferritin at 3.26 Å.

